# Using mixture density networks to emulate a stochastic within-host model of *Francisella tularensis* infection

**DOI:** 10.1101/2023.06.15.545189

**Authors:** Jonathan Carruthers, Thomas Finnie

**Affiliations:** Data, Analytics and Surveillance; UK Health Security Agency, Porton Down, UK

## Abstract

For stochastic models with large numbers of states, analytical techniques are often impractical, and simulations time-consuming and computationally demanding. This limitation can hinder the practical implementation of such models. In this study, we demonstrate how neural networks can be used to develop emulators for two outputs of a stochastic within-host model of *Francisella tularensis* infection: the dose-dependent probability of illness and the incubation period. Once the emulators are constructed, we employ Markov Chain Monte Carlo sampling methods to parameterize the within-host model using records of human infection. This inference is only possible through the use of a mixture density network to emulate the incubation period, providing accurate approximations of the corresponding probability distribution. Notably, these estimates improve upon previous approaches that relied on bacteria counts from the lungs of macaques. Our findings reveal a 50% infectious dose of approximately 10 colony-forming units and we estimate that the incubation period can last for up to 11 days following low dose exposure.

**Author summary:** *Francisella tularensis* is a highly infectious bacterium that remains in the top category of biothreat agents. Release of aerosolized bacteria could lead to many cases of acute and severe pneumonia over the days following. Mathematical modelling can contribute to the response to such an outbreak, combining dispersion models and disease models to identify the source of release and predict where cases are most likely to occur. However, these models can be computationally demanding and time consuming to run. In this article, we use neural networks to emulate the likelihood of disease and the duration of the incubation period from a stochastic within-host model. This enables rapid predictions to be made across a wide range of doses, thereby improving the practical applications of the model.

## Introduction

Mathematical modelling is an effective tool for describing the relationship between the quantity of biological agent an individual is exposed to and the likelihood that they develop disease [1]. Following the release of an agent, these models can provide critical insights into the source of the release and optimal prophylactic measures [2, 3].

Empirical dose-response models, such as the exponential and beta-Poisson models, rely on few parameters and are therefore quick and simple to fit. However, their predictions can vary hugely when extrapolating beyond the range of doses used for fitting [4]. This limitation is particularly pronounced when considering low dose exposure, where data is often sparse or non-existent and where it would be impractical to obtain even with direct experimentation. By contrast, mechanistic models build upon the underlying biological processes such that greater confidence can be placed on their predictions, even at these lower doses [5].

In a previous study, we developed a stochastic dose and time response model for human *Francisella tularensis* infection [6]. In conjunction with a zonal ventilation model, we demonstrated how it could be used to estimate the number of casualties in a microbiology facility following the accidental release of *F. tularensis* bacteria. One limitation of this model was that parameter inference was based on a comparison of model predictions with bacterial counts from the lungs of macaques, not humans. Although subsequent analysis showed that the estimates obtained were capable of explaining the incubation periods of human infection, the predicted credible regions at low doses were considerably wider than those observed. Furthermore, individual parameters were not identifiable from bacteria counts alone. Collectively, these concerns raise doubts about the reliability of the previous parameter estimates, and thus the wider application of the model. Ideally, the parameter inference should have been performed using the human infection data, rather than reserving it for validation purposes. However, due to time constraints in simulating the model, this was not possible. Since the existing model is stochastic, multiple realizations are necessary to fully understand the variability around quantities such as the incubation period. In addition, to account for heterogeneity within a population, certain model parameters were assumed to be random variables and were therefore capable of taking on multiple values. The need to sample across these possible values further increases the number of realizations required.

The time-consuming nature of simulating large stochastic models is a well-known limitation, particularly when using the exact Gillespie algorithm. As the population size increases, the time-step of the simulations decreases, leading to longer simulation times [7]. Approximate tau-leaping procedures overcome this issue by assuming that the rates of reaction change by such small amounts in a single time-step that they can be deemed constant, enabling multiple reactions to occur simultaneously [8]. Extensions of these methods have also been developed to further reduce simulation times, however, they often do so by exploiting the structure of the stochastic process and are therefore only applicable in certain scenarios. For example, a separate algorithm for stiff systems can be applied when reactions occur on different timescales, as is the case when modelling evolution within a growing virus population [9]. Rather than aiming to reduce the time required to simulate a model, an alternative approach is to construct an emulator [10, 11]. The use of emulators involves fitting a surrogate model to a set of existing simulations, which can then be used to rapidly predict the output of the original model [12, 13]. As a result, Monte Carlo inference methods and global sensitivity analyses, that require the model to be evaluated across many sets of inputs, can be applied more effectively.

In more recent work, this surrogate model takes the form of a mixture density network (MDN), a combination of a neural network and a mixture model [14]. In doing so, the outputs of the emulator specify a probability distribution rather than a single value. This is particularly useful when emulating stochastic models as it preserves the inherent variability of the system, avoiding the need to rely solely on summary statistics such as the mean. MDNs have successfully been implemented into a framework for emulating protein numbers in biochemical reaction networks, as well as emulating the infection dynamics of stochastic models of epidemics. [15, 16]. In both of these cases, however, the focus is on demonstrating the application of MDNs, so the baseline models have already been studied and parameterised in the literature. In this paper, our aim is provide a more practical application of MDNs to emulate a model. We begin by training an MDN to emulate the distribution of incubation periods from the within-host model of *F. tularensis* infection. By also creating an emulator for the probability of illness, we are then able to re-parameterise the original model using human infection data, a step that was previously not possible. The resulting model is the first mechanistic model to fully utilize this data and holds the potential to offer more reliable guidance for public health interventions in the event of a release of *F. tularensis* bacteria.

## Materials and methods

### Within-host model

The stochastic within-host model developed previously replicates the dynamics of *F. tularensis* in the lung [6]. Following the inhalation of some initial dose, bacteria can be taken up by host phagocytes with rate *α h*^*−* 1^ or cleared with rate *μ h*^*−* 1^. Infected phagocytes survive on average for 1/*δ* hours before rupturing and releasing their bacterial contents. The number of bacteria released on rupture is a random variable whose distribution is determined from a single-cell model describing the intracellular lifecycle of *F. tularensis*. This single-cell model was previously parameterised using *in vitro* measurements of intracellular bacterial growth and infected cell viability. Given that the recorded incubation periods from human infection used here are unlikely to provide any further insights into the intracellular dynamics, we continue to use the existing estimates of *δ* and the rupture distribution.

If all bacteria are successfully cleared from the lung then the individual recovers without any signs of illness. However, if the bacterial population grows to some threshold, *M*, then this is assumed to correspond to symptom onset. For a set initial dose, we are therefore interested in calculating the probability of reaching state *M*, as well as the time taken to reach state *M* . Given that a large number of bacteria is required for symptom onset, analytic methods have limited application and the model is instead simulated using a tau-leaping procedure with adaptive step size [8]. A single realization culminates in either illness or the elimination of bacteria, so multiple realizations are required to estimate the probability of illness and construct a distribution of incubation times. Together, these outputs allow us to specify the dose-response relationship and understand the variability in dose-dependent incubation times.

### Emulating the incubation period

A mixture density network (MDN) is used to emulate the distribution of incubation periods, a depiction of which is provided in Fig 1. Inputs from the within-host model are propagated through a series of dense layers, producing three sets of outputs. These outputs specify the means, standard deviations and weights of the Gaussian distributions that make up the mixture model. Here, we assume that a mixture of ten Gaussian distributions is sufficient to approximate the distribution of incubation periods for each unique set of inputs. To construct the MDN, we make use of the python interface Keras along with the Keras MDN Layer [17]. For each of the dense layers, we apply the commonly used rectified linear activation function (ReLU) since it introduces non-linearity but remains computationally efficient. For the nodes in the output layer that correspond to the standard deviations, we apply an exponential linear activation function offset by one to ensure that the standard deviations remain non-negative [18]. No activation functions are applied to the nodes representing the means or the weights,resulting in the weights being trained in logit space. To obtain probabilities that sum to one, a softmax function is subsequently applied to the final output weights.

**Fig 1.**
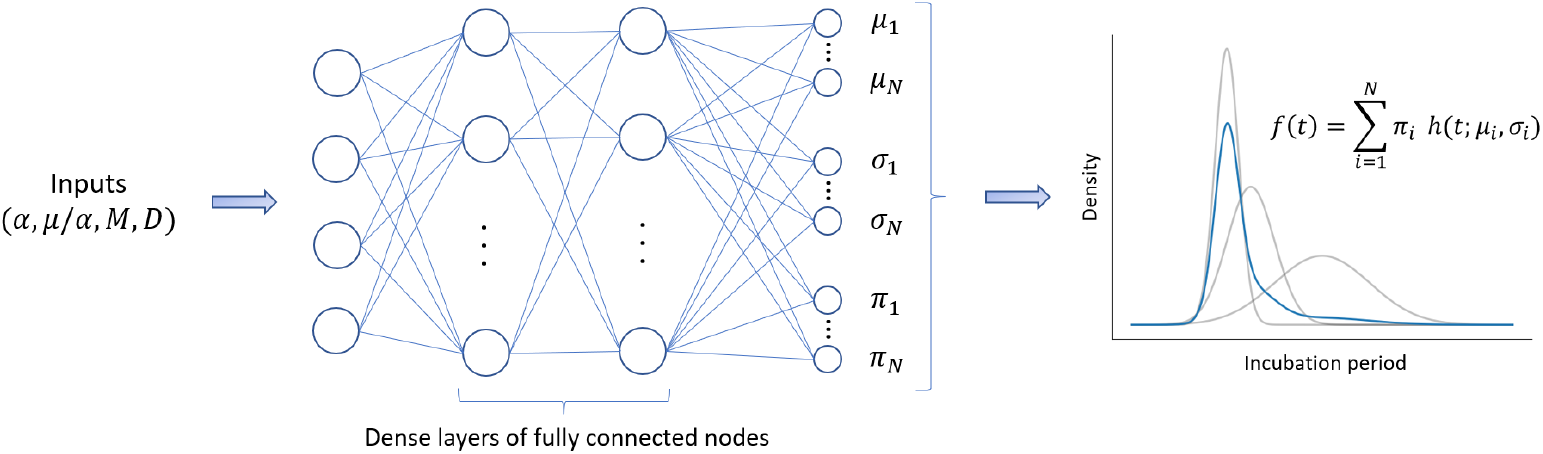
Mixture density network structure. Inputs from the within-host model are propagated through layers of fully connected nodes to estimate the means, standard deviations and weights for the individual components of the Gaussian mixture model.

With regards to the inputs, we previously demonstrated that there exists a strong correlation between the parameters *α* and *μ* [6]. This correlation is intuitive since *α* drives the growth of the bacterial population, whilst *μ* acts as a limiting factor.

Consequently, the ratio *μ/α* holds greater significance than the individual parameters themselves, so we include this as an input to the MDN. The remaining inputs consist of *α, M* and the initial dose, *D*. All four parameters are specified on a logarithmic scale and are normalized within the range [0,1]. This normalization facilitates the creation of a smooth loss surface and expedites the training process of the MDN. As the emulator’s ultimate purpose is to parameterise the within-host model, it is important to train it on a wide range of parameter values. The specific parameter ranges used in training are detailed in Table 1.

**Table 1.** Within-host model parameters.

| Parameter | Description | Emulator training bounds | Posterior |
| --- | --- | --- | --- |
| $\alpha$ | Rate of phagocytosis | $\log_{10} \alpha \sim U(-3, 0)$ | $7.4 \times 10^{-3} h^{-1}$ |
| $\mu/\alpha$ | Ratio of clearance and phagocytosis rates | $\log_{10}(\mu/\alpha) \sim U(-1.6, 1.6)$ | 14.26 |
| $D$ | Initial dose of inhaled bacteria | $\log_{10} D \sim U(0, 5)$ | — |
| $M$ | Threshold of bacteria required for symptom onset | $\log_{10} M \sim U(5, 20)$ | $\mu_M = 2.78 \times 10^8$ bacteria |

In order to train the MDN, 5 × 10^6^ unique parameter sets were sampled from the input space. For each parameter set, the within-host model was simulated using tau-leaping until a single incubation period was obtained. This means the MDN only observes a single sample from the distribution of incubation times, rather than the complete distribution. Given that these distributions are skewed, the incubation times were also converted to log-hours as this increases the chance of successfully approximating the full distribution with the mixture model. To fit the MDN, a stochastic gradient descent algorithm was employed, using the negative log-likelihood of the Gaussian mixture model as the loss function. Since the training sets contained only one incubation period, reserving a subset of these for testing purposes would not effectively assess the ability of the MDN to approximate the entire distribution.

Therefore, we instead generated a separate test set comprising 3 × 10^3^ inputs, with a sample of 10^3^ incubation times for each input. The Kolmogorov-Smirnov (KS) test statistic was initially used to measure the predictive capability of the MDN by comparing the cumulative density function of the mixture model with the empirical function specified by the test set samples [16]. A score close to zero would indicate a high level of agreement between the predicted and true distributions, whilst a score close to one would suggest little overlap between the distributions.

To determine an optimal architecture for the MDN, a grid search was performed. The number of dense layers ranged from 2 to 5, while the number of nodes in each dense layer was set to 64, 128, 256 or 512. To mitigate overfitting, the L2 regularization constant was also varied, resulting in training 600 different models. The best performing model was then selected as the one that minimised the proportion of test set predictions with a KS test statistic greater than 0.1. Following the grid search, approximately 83% of KS test statistics for the best performing model were below 0.1, with 39% falling below the critical value of 0.043 required for a one-sided KS test at a 5% significance level. Notably, higher KS test statistics were observed in regions of the input space where the initial dose of bacteria was largest. In such cases, the growth of bacteria is more deterministic, resulting in reduced variability in the time required for the population to reach *M* bacteria and hence a narrower range of incubation periods.

Comparisons between the predicted densities and the test set samples from the true distributions indicated that, in these scenarios, the MDN slightly underestimated or overestimated the mean. Given that the distributions are narrow, these deviations in the mean were amplified when calculating the KS test statistic. However, when converting the incubation times back from log-hours to days, the differences were minor and within an acceptable degree of accuracy for the intended purpose of the model.

With this in mind, and to ensure the optimal model reflects our needs, a second grid search was performed using the following scaled KS distance

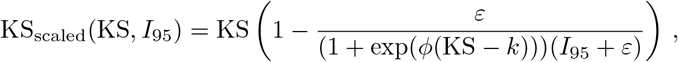

where KS is the original test statistic and *I*_95_ is the inter-95 percentile range. This scaled distance reduces the penalty when the range of incubation periods is narrow but the model prediction yields a larger KS test statistic. When KS *> k*, KS_scaled_ KS and no scaling is applied. Setting *k* = 0.8 ensures that predictions that have little to no overlap with the true distribution retain a high distance and thus a model that produces such predictions would be less favoured. The right-hand panel of Fig 2 demonstrates that this distance results in a more uniform scoring for better performing models.

**Fig 2.**
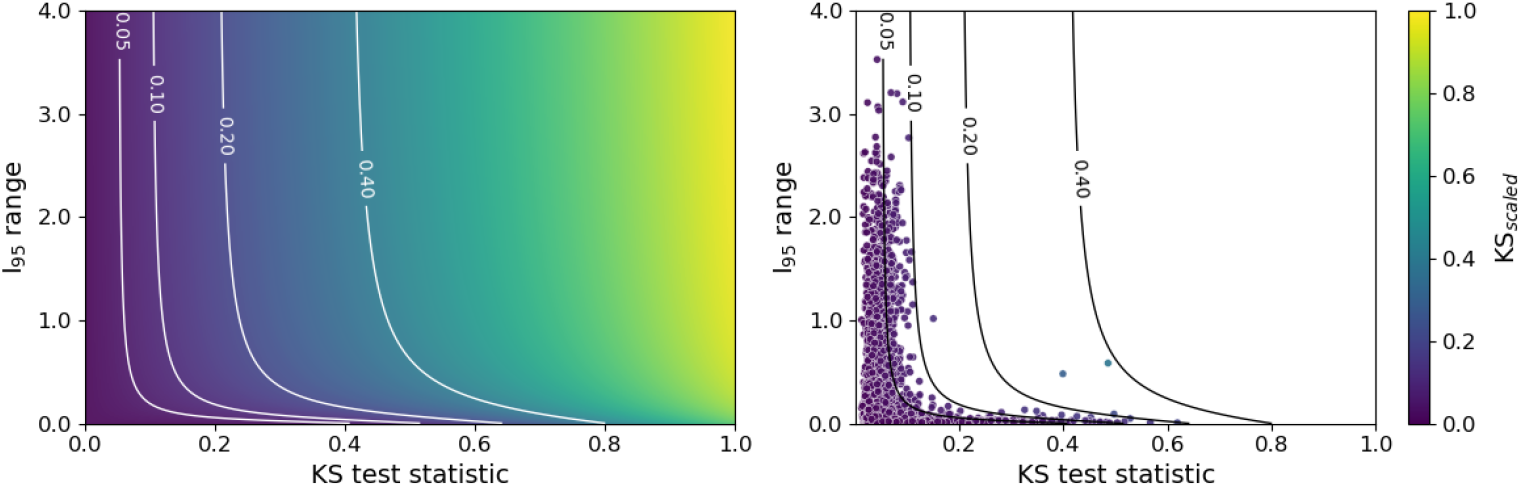
Scaled Kolmogorov-Smirnov distance. Left: KS_scaled_ as a function of the original KS test statistic and the inter-95 percentile range. Right: KS test statistics and inter-95 percentile ranges from an initial grid search across model architectures; points are shaded by the corresponding value of KS_scaled_. Here, *ε*=0.2, *k*=0.8 and *ϕ*=5.

### Emulating the probability of illness

An emulator for the probability of illness can be created using a fully-connected neural network with an output layer consisting of just a single node. Whilst the full within-host model must be considered to evaluate the incubation period, the state space can be truncated to a ‘point of no return’ when calculating the probability of illness [6, 19]. This truncation represents a state where the probability of elimination is approximately zero and, if reached, the stochastic process is almost guaranteed to continue to state *M* . As a result, not only can *M* now be ignored, reducing the dimensionality of the input space, but it is also no longer necessary to include such high initial doses. For this model it is suitable to truncate the state space such that states corresponding to more than 500 bacteria are removed.

By truncating the state space, matrix analytic methods can be used to determine the probability of illness, as described in [6]. Additionally, this approach provides a more comprehensive training set by yielding the probability of illness across all doses for a single (*α, μ/α*) pair. To construct a training set, we therefore sample 500 (*α, μ/α*) pairs from the ranges provided in Table 1. The probability of illness is then evaluated for initial doses ranging from 1 to 250 bacteria, effectively generating a complete dose-response curve for each pair. When the ratio *μ/α* is less than 1, the clearance rate of bacteria is lower than the rate at which they infect cells, resulting in a probability of illness close to 1 even at low initial doses. Consequently, out of the 1.25 × 10^5^ training sets, 78% of them have a probability of illness equal to 1. Recognizing that much of this training data contributes little additional information to the emulator, we choose to thin the doses once the probability reaches 1, retaining approximately 5 doses for each (*α, μ/α*). In doing so, the final training set consists of 8.5 10^3^ inputs and corresponding outputs.

As with the MDN, a grid search is performed across different model architectures, with the number of dense layers ranging from 2 to 5 and the number of nodes in each dense layer equal to 64, 128 or 256. A ReLU activation function is also applied to nodes in each dense layer, whilst a sigmoidal activation function is used for the output layer to ensure the network returns a probability. We fit the neural network to the training set using a stochastic gradient descent algorithm, where the loss function is given by the mean squared error. The best performing model from the grid search is selected as the one that maximises the the fraction of predictions that are within a 5 × 10^*−* 3^ error of the true probability, for both the training set and 10^3^ separate test sets.

### Parameter inference

Let ***D*** = (*D*_1_, …, *D*_*n*_) and ***x*** = (*x*_1_, …, *x*_*n*_) respectively denote the doses of bacteria that a set of individuals inhale, and whether or not they developed illness. Here, *x*_*i*_ = 1 corresponds to illness, with *x*_*i*_ = 0 otherwise. The likelihood of the *n* observed outcomes is given by

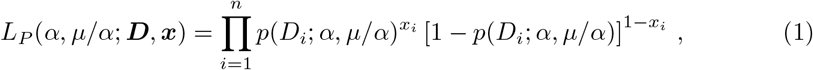

where *p* is the dose-dependent probability of illness. Now let ***t*** = (*t*_1_, …, *t*_*n*_) be the number of days between exposure and symptom onset for those individuals that developed illness. Here, the observed incubation periods are interval censored given that only the day of symptom onset is provided. As a result we consider the likelihood of symptom onset occurring across the 24-hour period that precedes the recorded values, noting that predictions from the MDN are in log-hours. At high doses the variability in predicted incubation times is too small to explain the variability in the observed incubation periods. Previously, Wood *et al*. assumed that the threshold required for symptom onset was not fixed but instead followed a log-normal distribution, representing heterogeneity across individuals and resulting in greater variability in predicted incubation periods [19]. However, their specific parameterisation of the log-normal distribution suggests that *M* could be as high as 10^16^ CFUs. Here, we continue to assume that log_10_ *M* ∼ 𝒩 (*μ*_*M*_, *σ*_*M*_), but set *σ*_*M*_ = 1 and infer *μ*_*M*_ .

Collectively, these assumptions yield the following likelihood function for the incubation periods

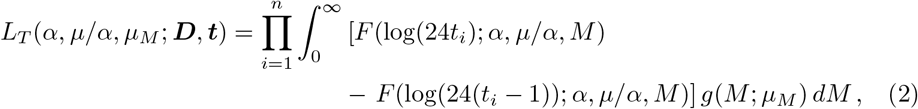

where *F* is the cumulative density function of the dose-dependent incubation period, evaluated using the MDN, and *g* is the density of a Gaussian distribution with mean *μ*_*M*_ and unit variance. Since the probability of illness and the incubation period are independent, the combined likelihood is the product of the individual likelihoods, that is,

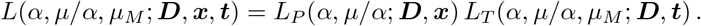

Posterior probability likelihood distributions are obtained using the python MCMC implementation emcee with the above likelihood [20]. Both *α* and *μ/α* are assumed to follow a uniform prior distribution with ranges equal to those used in the training of the emulators. For *μ*_*M*_, a *U* (8.25, 16.75) prior is used since this ensures that the distribution of log_10_ *M* is contained within the bounds used to train the MDN.

## Results

For the distribution of incubation periods, the best performing MDN architecture consists of 3 dense layers, each comprising 512 nodes. The top-left panel of Fig 3 shows a histogram of KS_scaled_ values for this model, with 94% of the 3 × 10^3^ test sets resulting in a value below 0.05. From Fig 2, we can see that these values would remain small when considering the original KS test-statistic, even for those incubation periods with narrow ranges. This suggests that the MDN is able to accurately capture the variability in incubation periods. As an example of the MDN output, the bottom panels of Fig 3 show the predicted and true distributions for a single set of inputs, alongside the components of the mixture model that have a weight greater than 10^*−* 2^. The MDN never utilizes the full set of available Gaussian distributions, instead finding that two or three components are sufficient to describe the target distribution for the majority of test sets. For the remaining components, the weights are approximately zero.

**Fig 3.**
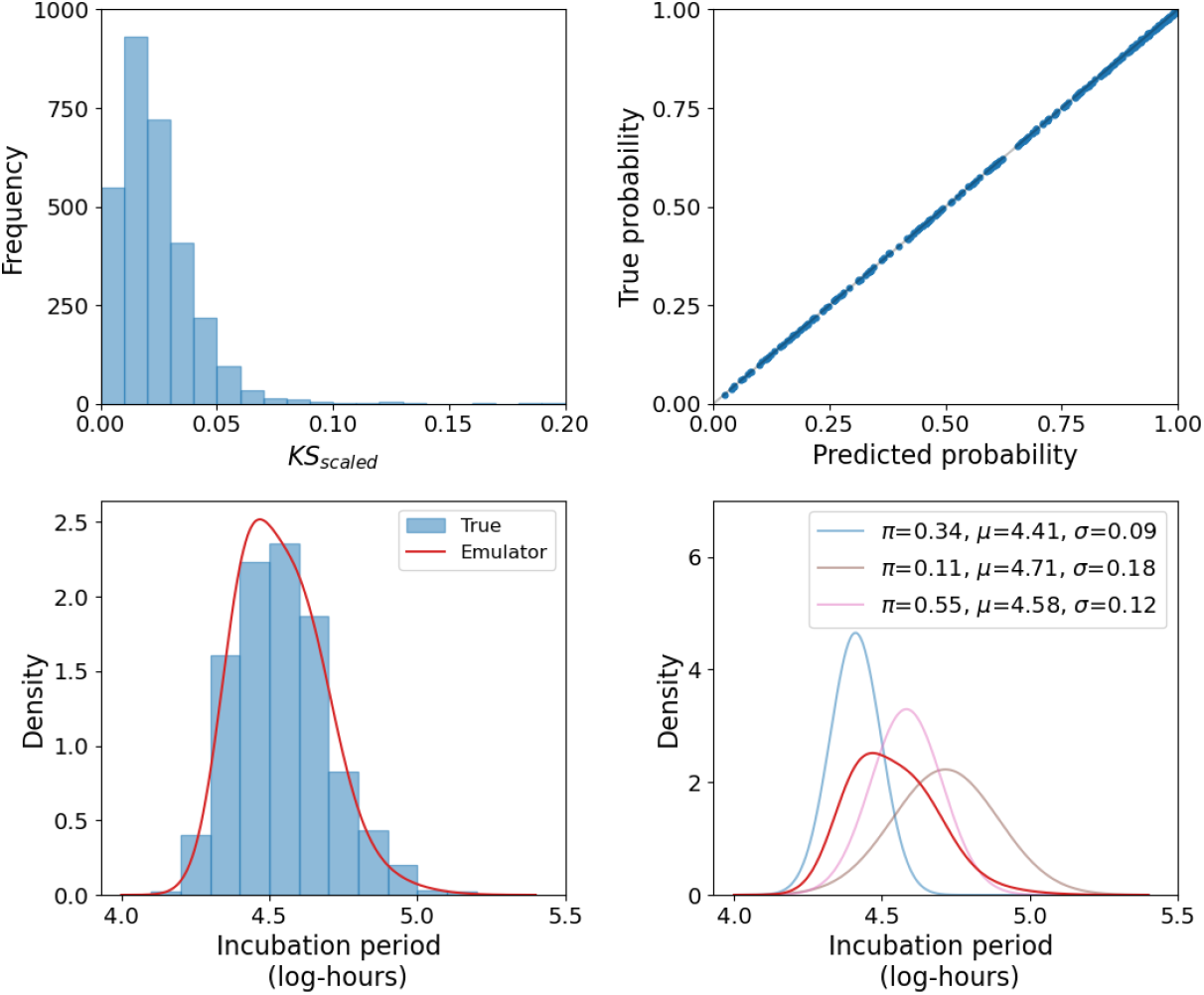
Summary of emulator performance. Top left: Histogram of KS_scaled_ scores for the best performing MDN. Top right: Comparison between the true and predicted probability of illness, showing near perfect agreement. Bottom: For a single set of inputs, a comparison between the distribution predicted by the MDN and the true distribution from stochastic simulations. For the same example, the primary components of the mixture distribution are also shown.

The neural network that most accurately emulates the probability of illness has 3 dense layers, each containing 128 nodes. The top-right panel of Fig 3 compares the true probability from the unseen test set with predictions from the emulator, demonstrating that it is able to predict the probability with near-perfect accuracy. This high level of precision is understandable given the relatively low dimensionality of the input space. Furthermore, the probability of illness is most sensitive to *μ/α* and the dose. Varying the dose shifts the output along a dose-response curve, a smooth sigmoidal function.

For large values of *μ/α*, indicating a greater level of bacterial clearance, this dose-response curve is centred around higher doses, whilst reducing *μ/α* gradually shifts it towards lower doses. The smooth nature of this output space enables the emulator to make more accurate predictions about untrained regions. With accurate emulators that significantly reduce the computational time required to evaluate the incubation period and probability of illness, it is now possible to apply them to the process of parameter inference.

Beginning in the 1950s, Operation Whitecoat was a vaccine research program investigating the human response to potential biothreat agents, including *F. tularensis* [21]. During this program, human volunteers were exposed by inhalation to the highly infectious Schu S4 strain and observed for symptoms associated with pneumonic tularemia [22]. Estimated inhaled doses varied from as little as 10 CFUs to 6.2×10^4^ CFUs. For the 118 individuals that were exposed to *F. tularensis*, McClellan *et al*. provides records of whether or not they developed febrile illness. At doses above 46 CFUs, febrile illness was observed in all individuals, suggesting that the majority of the data-set is redundant when considering the dose-response relationship. As a result, when constructing the likelihood in Eq (1), only the outcomes for the 22 volunteers who were exposed to doses below 100 CFUs are used. For two subsets of individuals, one exposed to low doses (10-52 CFUs) and the other receiving high doses (2.5 × 10^4^ CFUs), the number of days between exposure and onset of illness are also provided [23, 24]. The incubation times for these 32 volunteers are used to construct the likelihood in Eq (2).

An MCMC algorithm is used to sample from posterior distributions for *α, μ/α* and *μ*_*M*_, the mean threshold of bacteria required for symptom onset. Posterior probability likelihood distributions for each of these parameters are provided in Fig 4. These narrow distributions, together with low correlations between parameters, suggests that all three parameters are identifiable. We estimate that *F. tularensis* bacteria are cleared from the lungs of individuals at rate *μ* = 0.1 *h*^*−* 1^ while the uptake and survival of bacteria by host phagocytes occurs at rate *α* = 7.4 × 10^*−* 3^ *h*^*−* 1^. The distribution of *μ*_*M*_ is skewed towards lower values of the prior distribution, however, reducing the lower bound of the prior distribution is not possible as this would allow *M* to extend to values beyond those used to train the emulator. Although additional training could be conducted, we believe that the current lower bound of 10^5^ bacteria is sufficiently small given that it represents the number of bacteria associated with symptom onset. To assess the sensitivity of the inference to fixing *σ*_*M*_ = 1, it has been repeated for *σ*_*M*_ = 0.75 and *σ*_*M*_ = 1.25 (Fig 5). The resulting samples show little differences in *α* and *μ/α*, suggesting that these are robust to changes in *σ*_*M*_ . For *μ*_*M*_, however, larger values correspond to larger *σ*_*M*_, with posterior medians of 4.64 × 10^7^ and 1.18 × 10^9^ bacteria for *σ*_*M*_ = 0.75 and *σ*_*M*_ = 1.25, respectively.

**Fig 4.**
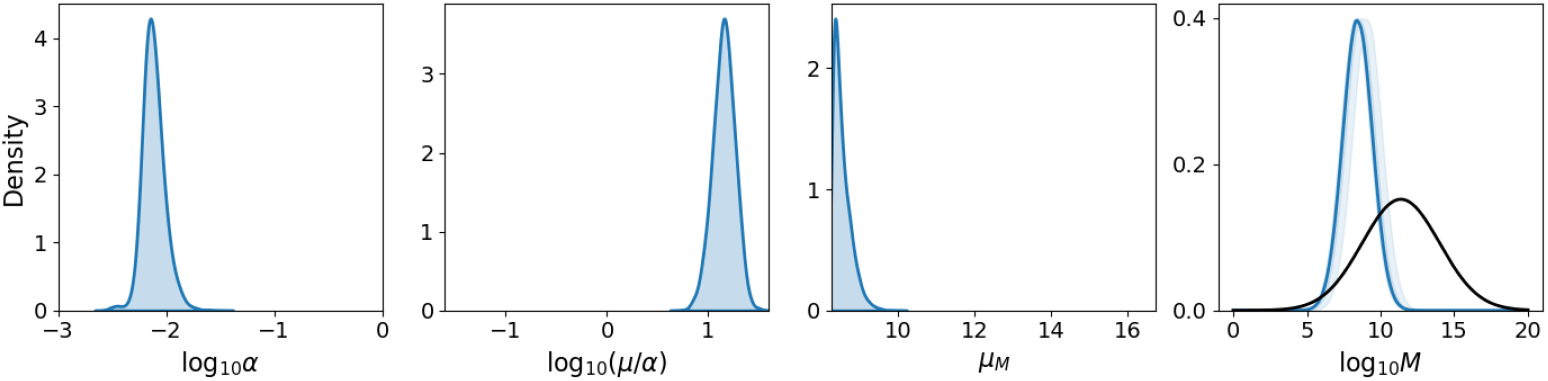
Posterior probability likelihood distributions. Distributions for within-host parameters are inferred from the Operation Whitecoat data [21, 23, 24]. The right-hand plot shows the resulting distribution of *M*, the threshold of bacteria required for symptoms, compared to the existing distribution proposed by Wood *et al*. (black) [19].

**Fig 5.**
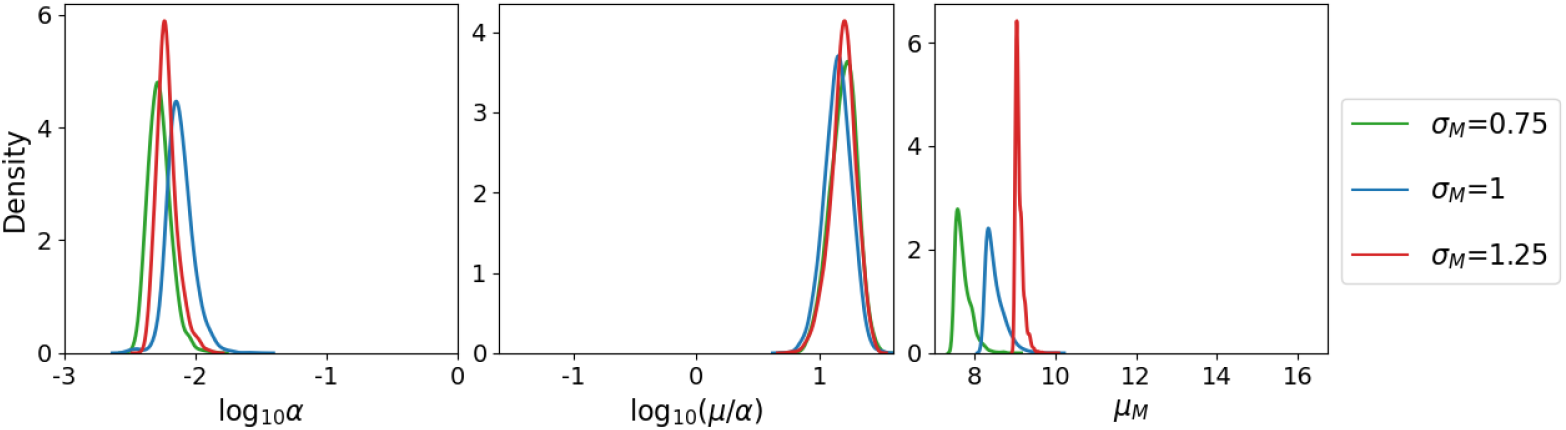
Posterior probability likelihood distributions for varying. *σ*_*M*_ . Distributions obtained after repeating the inference for varying values of *σ*_*M*_, the standard deviation of the threshold log_10_ *M* .

Fig 6 provides posterior predictions for the probability of illness and incubation period as a function of dose. From this we estimate that the ID_50_, the dose required to produce infection in 50% of individuals, is approximately 10 CFUs. For the incubation period, the majority of observed times fall within the middle 50% of the predicted distribution, with almost all lying within the middle 95%. The only exception to this is the one individual who, despite being exposed to a high initial dose, did not develop symptoms until a week later. As a comparison, the 95% confidence regions from a statistical model by McClellan *et al*. are also provided [21]. In this model, the logarithm of the incubation time is assumed to follow a Gaussian distribution with constant variance and a mean that is a decreasing function of dose. The data used to fit the model also contain more precise estimates of the incubation periods that were derived from temperature profiles of each volunteer. Whilst the predictions of the two models are similar, our mechanistic model predicts a notable reduction in variability once the dose exceeds 100 CFUs, which is not captured by the statistical model. The reason for this reduction is that at high doses, much of the variability in incubation periods is due to variability in the threshold, *M* [19]. The credible regions are therefore of equal width because the distribution of *M* remains the same across all doses. In contrast, at lower doses, stochastic effects contribute more to the overall variability. If the early growth of the bacterial population is slower, it takes longer to reach the threshold of *M* bacteria, extending the duration of the incubation period. From Fig 6, it appears that the importance of these stochastic effects significantly diminishes once the dose exceeds 100 CFUs.

**Fig 6.**
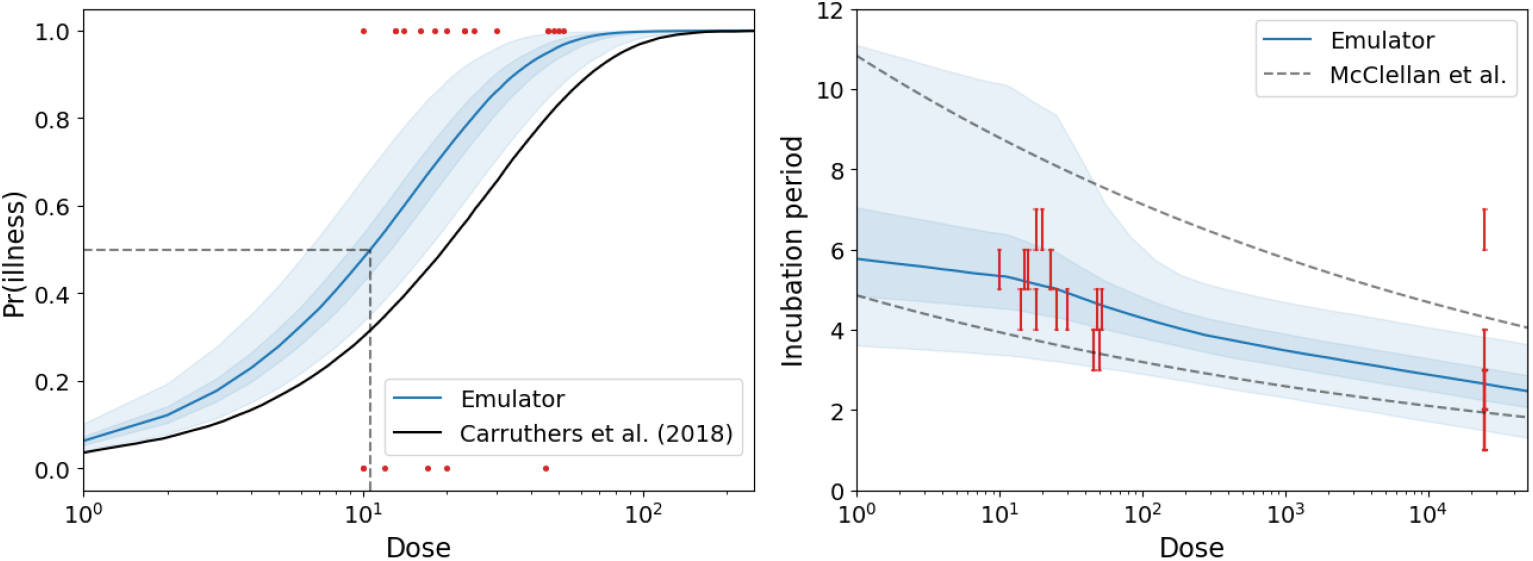
Posterior predictions. Left: Emulator predictions for the probability of illness. Shaded regions represent 50% and 95% credible regions, the dashed line indicates the median infectious dose (ID_50_) and the solid black line shows the previous dose-response curve. Right: Emulator predictions for the incubation period, the dashed line corresponds to 95% confidence regions predicted by McClellan *et al*. [21]. Red bars correspond to the 24-hour period during which symptom onset occurred.

## Discussion

In this paper, we have demonstrated how neural networks can be used to create emulators for key descriptors of a stochastic within-host model of *F. tularensis* infection, namely the probability of illness and the distribution of incubation periods. Reducing the computational time required to evaluate these has allowed us to combine more powerful methods for parameter inference with data-sets that better reflect human infection.

Given the difficulty in constructing the likelihood for stochastic mechanistic models, we previously relied on approximate Bayesian computation to perform the parameter inference, a likelihood-free method [6]. Whilst this is commonly used, the original approach is an inefficient way to sample from the posterior distribution and the success of more recent sequential approaches can depend on the choice of tolerances [25, 26]. Using an MDN to approximate the distribution of incubation times by a mixture of Gaussian distributions provides us with a simple expression for the cumulative density function that can be used to specify the likelihood. Together with the reduced computational time, this allowed us to apply an MCMC sampling method whilst also incorporating variability in the threshold, *M* . Such an approach would not have been possible when relying solely on simulations of the stochastic model.

From the results of the previous parameter inference, that relied on bacterial counts from the lungs of macaques, posterior median estimates of *α* and *μ* were larger than those obtained here [6]. However, the strong correlation between these parameters meant that the estimates were heavily influenced by the choice of prior distribution. When comparing the ratio *μ/α*, our posterior median estimate of 14.3 is also lower than the previous estimate of 25.9. Smaller values of *μ/α* indicate a reduced resistance to bacterial growth from clearance mechanisms, increasing the likelihood of reaching the threshold required for illness. This suggests that we previously underestimated the probability of illness at low dose exposures, as shown in the left-hand panel of Fig 6. As a result, the estimated ID_50_ has almost halved, from 19 CFUs to 10 CFUs. This new estimate is more reliable given it is inferred directly from the outcome of human infection, rather than from *in vivo* measurements of bacterial counts.

We estimate that a single bacterium survives phagocytosis with probability *α/*(*μ* + *α*) = 6.6 × 10^*−* 2^. This is 10-times smaller than an equivalent estimate obtained by Heppell *et al*., who adapt a mechanistic model of *Coxiella burnetii* infection to describe *F. tularensis* infection [27]. This difference can be attributed to their assumption that not all bacteria are deposited in the lung. A key feature of their model is the inclusion of a 5-lobe lung model to estimate the fraction of bacteria deposited in the pulmonary region of the lung, accounting for the particle size distribution of the aerosol and the breathing rate of adults. They estimate that only 11% of inhaled bacteria will ultimately be deposited, effectively reducing the initial dose. Treating this as the instantaneous clearance of bacteria, it is understandable that we estimate a higher proportion of bacteria being cleared, since this immediate loss is instead averaged across the initial course of infection.

One of the reasons for repeating the parameter inference was to more accurately estimate the distribution of the threshold, *M* . The existing log-normal distribution initially proposed by Wood *et al*. allows this threshold to exceed 10^16^ CFUs, as shown in the right-hand panel of Fig 4 [19]. Our new estimate reveals a narrower distribution with a lower mean, suggesting that the threshold rarely exceeds 10^11^ bacteria. This is more comparable to values of 7.9 × 10^6^ and 1.6 × 10^8^ CFUs identified in existing modelling studies, and represents a value that is more aligned with the realistic volume of the human lung [27, 28]. Note that these previous studies do not offer any insights into the variability of the threshold. Whilst the distribution obtained here represents an improvement over the previous distribution, some uncertainty remains. When repeating the inference for different values of *σ*_*M*_, we observed a positive correlation between *σ*_*M*_ and the estimate of *μ*_*M*_, with larger values yielding slightly higher likelihoods. Despite this, larger values of *σ*_*M*_ and *μ*_*M*_ give rise to distributions similar to those predicted by Wood *et al*., which we believe to be implausible from a biological perspective.

Ultimately, this threshold is a surrogate for inflammation driven by the host immune response and specifying an exact value will always remain a challenge. As long as the threshold is large enough, its specific value does not affect the probability of illness, only the duration of the incubation period. Whilst other biological markers may correlate better with symptom onset, incorporating them would increase model complexity, and similar issues would arise without accurate measurements. Specifying an appropriate marker and determining its value is a key challenge we face when developing mechanistic dose and time response models.

With a fully parameterised within-host model of human *F. tularensis* infection and emulators that enable efficient evaluation of its outputs, future work can focus on integrating this model into practical applications. In the event of a release of *F. tularensis* bacteria, back-calculation techniques can be used to estimate the time and source of release from those individuals presenting symptoms [3]. These techniques combine dispersion models, which forecast the concentration of bacteria at specific locations, with a disease model that determines the likelihood of the observed cases occurring. Once the source has been identified it is possible to predict the size of the affected region. This is where knowledge of low dose exposure is particularly important, as this determines the boundaries of the region. Inaccuracies at this stage can result in miscalculating the number of cases and, consequently, the required quantity of antibiotics. In this context, the model presented here serves as a valuable disease model, providing dose-dependent probabilities of illness and incubation periods that can ultimately contribute to enhancing outbreak response strategies. Incorporating the underlying biological mechanisms also provides us with more reliable predictions at lower doses.

